# Information topology of gene expression profile in dopaminergic neurons

**DOI:** 10.1101/168740

**Authors:** Mónica Tapia Pacheco, Pierre Baudot, Martial A. Dufour, Christine Formisano-Tréziny, Simone Temporal, Manon Lasserre, Jean Gabert, Kazuto Kobayashi, Jean-Marc Goaillard

## Abstract

Extracting high-degree interactions and dependences between variables (pairs, triplets, … *k*-tuples) is a challenge posed by all omics approaches^**1, 2**^. Here we used multivariate mutual information (I_k_) analysis^**3**^ on single-cell retro-transcription quantitative PCR (sc-RTqPCR) data obtained from midbrain neurons to estimate the k-dimensional topology of their gene expression profiles. 41 mRNAs were quantified and statistical dependences in gene expression levels could be fully described for 21 genes: I_k_ analysis revealed a complex combinatorial structure including modules of pairs, triplets (up to 6-tuples) sharing strong positive, negative or zero I_k_, corresponding to co-varying, clustering and independent sets of genes, respectively. Therefore, I_k_ analysis simultaneously identified heterogeneity (negative I_k_) of the cell population under study and regulatory principles conserved across the population (homogeneity, positive I_k_). Moreover, maximum information paths enabled to determine the size and stability of such transcriptional modules. I_k_ analysis represents a new topological and statistical method of data analysis.

## MAIN TEXT

The recent evolution of single-cell transcriptomics has created much hope for our understanding of cell identity, cell development and gene regulation^**4**^. Using quantitative PCR or RNAseq, tens to thousands of mRNAs can be quantified from a single cell, generating particularly high-dimensional datasets (gene expression profiles). Combined with clustering and dimensionality-reduction techniques, these approaches have been successfully used to identify and separate cell types in various tissues, including the brain^**5**^. Single-cell transcriptomics has also be used to shed light on the gene regulatory principles underlying the specific phenotype of different cell types^6, 7^, frequently relying on pairwise analysis of gene expression levels to infer gene regulatory networks^6, 8^. However, the modular architecture of gene networks suggests that extracting higher-degree interactions between gene expression profiles may be necessary to understand gene regulation, and various approaches based on probability/information theory^8–10^ or homology^**11**^ have been proposed to tackle this issue.

Several transcriptomics studies have been performed on midbrain dopaminergic (DA) neurons^5, 12^: consistent with the heterogeneous vulnerability of this neuronal population in Parkinson’s disease^**13**^, qPCR and RNAseq performed at the single-cell level have revealed a significant diversity in gene expression profiles^5,12^. In parallel, much work has also been performed to understand the gene regulatory networks and identify the regulatory factors underlying the emergence of the DA phenotype^**14**^, with the therapeutical intent of producing functional DA neurons from induced pluripotent stem cells^**15**^.

Here we implement multivariate mutual information (I_k_) analysis on transcriptomics data from single midbrain DA neurons to simultaneously provide new insights about the molecular heterogeneity of this neuronal population and about the gene regulatory principles underlying its specific phenotype.

We performed sc-RTqPCR on acutely dissociated identified midbrain neurons using the microfluidic BioMark™ HD Fluidigm platform. TH-GFP mice were used to preferentially target putative DA neurons (identified by the presence of tyrosine hydroxylase, TH-positive neurons, **Supplementary Figure 1a**). Electrophysiological recordings confirmed that acutely dissociated GFP and non-GFP neurons displayed the electrical properties expected for DA and non-dopaminergic (nDA) midbrain neurons^16, 17^, respectively (**Supplementary Figure 2**). However, since TH presence alone has been shown to not be a reliable marker^**18**^, DA and nDA phenotypes were refined based on the combined expression of *Th*/TH and *Slc6a3*/DAT (DA transporter) or lack thereof, allowing neurons collected from wild-type animals to be included (**Supplementary Figure 1b**). Based on *Th*-*Slc6a3* expression, 111 neurons were classified as DA and 37 as nDA.

We quantified the levels of expression of 41 genes (**Figure 1a**), including 19 related to ion channel function, 9 related to neurotransmitter definition, 5 related to neuronal activation and calcium binding, and 3 related to neuronal structure (**Supplementary Figure 1c**). As expected, DA metabolism and signaling-related genes such as *Th*/TH, *Slc6a3*/DAT, *Slc18a2*/VMAT2, *Drd2*/D2R were highly expressed in DA neurons only, while expression levels of *Slc17a6*/VGLUT2, *Gad1*/GAD67 and *Gad2*/GAD65 suggested that collected nDA neurons used mainly glutamate or GABA as neurotransmitters (**Figure 1a-b**, **Supplementary Figure 3**). While some ion channels showed similar expression profiles in DA and nDA neurons (*Cacna1c*/Cav1.2, *Cacna1g*/Cav3.1, *Hcn2*/HCN2, *Hcn4*/HCN4, *Kcna2*/Kv1.2, *Scn8a*/Nav1.6), others (*Kcnb1*/Kv2.1, *Kcnd3_2*/Kv4.3, *Kcnj6*/GIRK2, *Kcnn3*/SK3, *Scn2a1*/Nav1.2) displayed higher levels of expression in DA neurons (**Figure 1b**, **Supplementary Figure 3**). In addition, although a few genes displayed a fairly stable level of expression across DA neurons (*Th*/TH, *Slc6a3*/DAT, *Kcnd3_2*/Kv4.3, *Scn2a1*/Nav1.2), most genes displayed significant variability in their expression levels (including dropout events) across cells (**Figure 1b**, **Supplementary Figure 3**), consistent with the already documented heterogeneity of midbrain DA neurons^**5, 13, 14**^.

**Figure 1.**
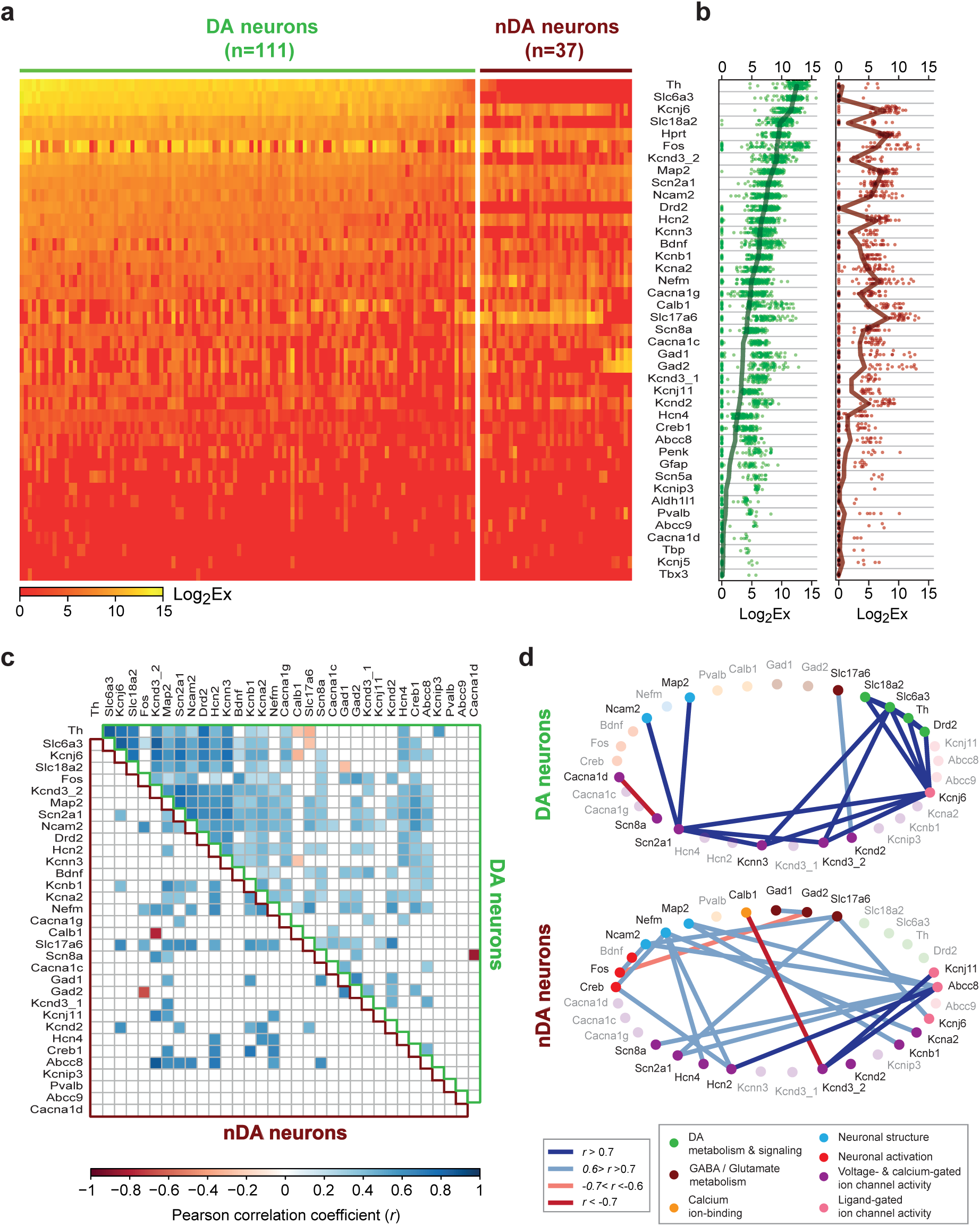
First and second order linear analysis reveals strong correlations in gene expression levels in midbrain DA and nDA neurons. **a**, heatmap representing the levels of expression of 41 genes in the collected 111 DA and 37 nDA neurons. Neurons are ordered based on *Th* and *Slc6a3* levels of expression, and genes are ordered based on their average level of expression in DA neurons (see **b**, left plot). **b**, levels of expression of the 41 genes presented in **a** in the DA population (left, green) and in the nDA population (right, dark red). The thick green and red lines represent the average expression levels while each dot corresponds to the expression level in one neuron. **c**, heatmap representing the significant correlations in expression levels for 33 genes in DA neurons (upper right triangle) and nDA neurons (lower left triangle) (Pearson correlation coefficient). Correlations were processed on non-zero values of expression, and only correlations with a p value < 0.05 and n > 5 are represented. Please note the difference in patterns of correlations between DA and nDA neurons. **d**, scaffold representations of the 20 most significant correlations in expression levels in DA (top) and nDA (bottom) neurons (r values > 0.6 or < −0.6, see color coding in the left box). mRNAs were ordered based on the known function of the corresponding proteins (see right box for the color coding of functions). The genes involved in the depicted correlations are highlighted (dark font, bright colors). Please note the strong connectivity between DA metabolism/signaling and ion channel genes in DA neurons.

As a first step in deciphering higher-degree relationships, we performed Pearson correlation analysis on the 33 most relevant genes (**Figure 1c-d, Supplementary Figure 4**). The patterns of correlations were clearly different for DA and nDA neurons, with more widespread correlations in DA neurons, as can be seen in the correlation maps (**Figure 1c**). This is only partly surprising as most of the genes were chosen because of their known expression in DA neurons, but it nonetheless demonstrates that specific signatures of second-degree linear relationships participate in the identity of the two populations under study (**Figure 1d**). While most of the cell type-specific correlations involved differentially expressed mRNAs, some similarly expressed genes displayed a stronger correlation in a specific population: *Kcnj6*/GIRK2 *vs Scn2a1*/Nav1.2 for instance in DA neurons, *Scn2a1*/Nav1.2 *vs Slc17a6*/VGLUT2 or *Hcn4*/HCN4 *vs Nefm/*NEF3 in nDA neurons (**Figure 1d**, **Supplementary Figure 4**). Several correlations were also present in both cell types (*Kcna2*/Kv1.2 *vs Nefm*/NEF3, *Hcn2*/HCN2 *vs Nefm*/NEF3). Interestingly, some of the strongest correlations found in DA neurons linked the group of genes involved in DA metabolism and signaling (*Th*/TH, *Slc6a3*/DAT, *Slc18a2*/VMAT2, *Drd2*/D2R) to a group of ion channel genes (*Kcnj6*/GIRK2, *Kcnd3_2*/Kv4.3, *Kcnn3*/SK3, *Scn2a1*/Nav1.2) (**Figure 1d**, **Supplementary Figure 4**), suggesting the existence of a large module of co-regulated genes. However, the size of such modules might only be accurately defined by methods capturing high-dimensional (beyond pairs) statistical dependences.

Various information theoretical approaches have been proposed to define gene regulatory modules based on the exploration of higher-degree relationships, notably three-way interactions^**8-10**^ (see also **Supplementary methods**). Here we present a method that combines in a single framework statistical and topological analysis of gene expression for systematic identification and quantification of such regulatory modules, based on the information cohomology developed by Baudot and Bennequin^**3**^. In this framework, joint-entropy (H_k_) and multivariate mutual information (I_k_) quantify the variability/randomness and the statistical dependences of the variables, respectively, while simultaneoustly estimating the topology of the dataset. We restricted the general setting defining information structures from the whole lattice of partitions of joint random variables to the simplicial sublattice of “set of subsets”, thus computationally allowing an exhaustive estimation of H_k_ and I_k_ at all degrees *k* and for every *k*-tuple (for *k* ≤ *n*=21, *k* being the degree/number of genes analyzed as a *k*-tuple, *n* being the total number of genes analyzed; **Figure 2a, Supplementary methods).** Information values obtained with this analysis provide a ranking of the lattices at each degree *k* (**Supplementary methods**). The H_k_ and I_k_ analysis therefore estimate the variability and statistical dependences at all degrees *k*, from 1 to *n*. I_k_ is defined as follows^**3, 19, 20**^:

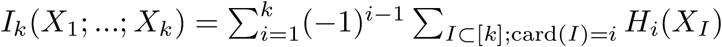

giving, for *k*=3,

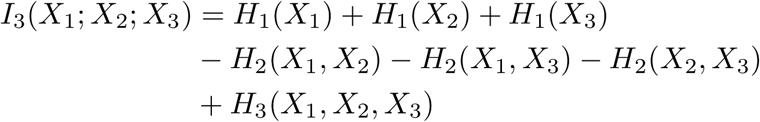

where *X_I_* denotes the joint-variable corresponding to the subset *I*. I_k_ is equivalent to entropy for *k* = 1, has upper and lower limit values of log_2_(*N*) and –log_2_(*N*) bits (*N* being the number of bins or graining used to discretize the data; *N*=8 in the present case, **Supplementary Figure 5**), is always non-negative for *k* < 3, and can take negative values for *k* ≥ 3 ^**19-21**^ (**Supplementary methods**). As an example, the maxima and minima of I_3_ for 3 binary variables are depicted in **Supplementary Figure 6**: while maxima (positive I_k_) correspond to a fully redundant behavior (x_1_, x_2_ and x_3_ are informationally equivalent), the minima (negative I_k_) correspond to cases where variables are pairwise independent (I_2_=0) but strictly tripletwise dependent (emergent behavior). In other terms, positive I_k_ captures co-variations and usual linear correlations as a subcase, zeros of I_k_ capture statistical independence, and negativity captures more complex relationships that cannot be detected on lower dimensional projections, such as degree-specific clustering patterns (also called synergy or frustation)^**9, 21, 22**^ (**Supplementary methods**).

**Figure 2.**
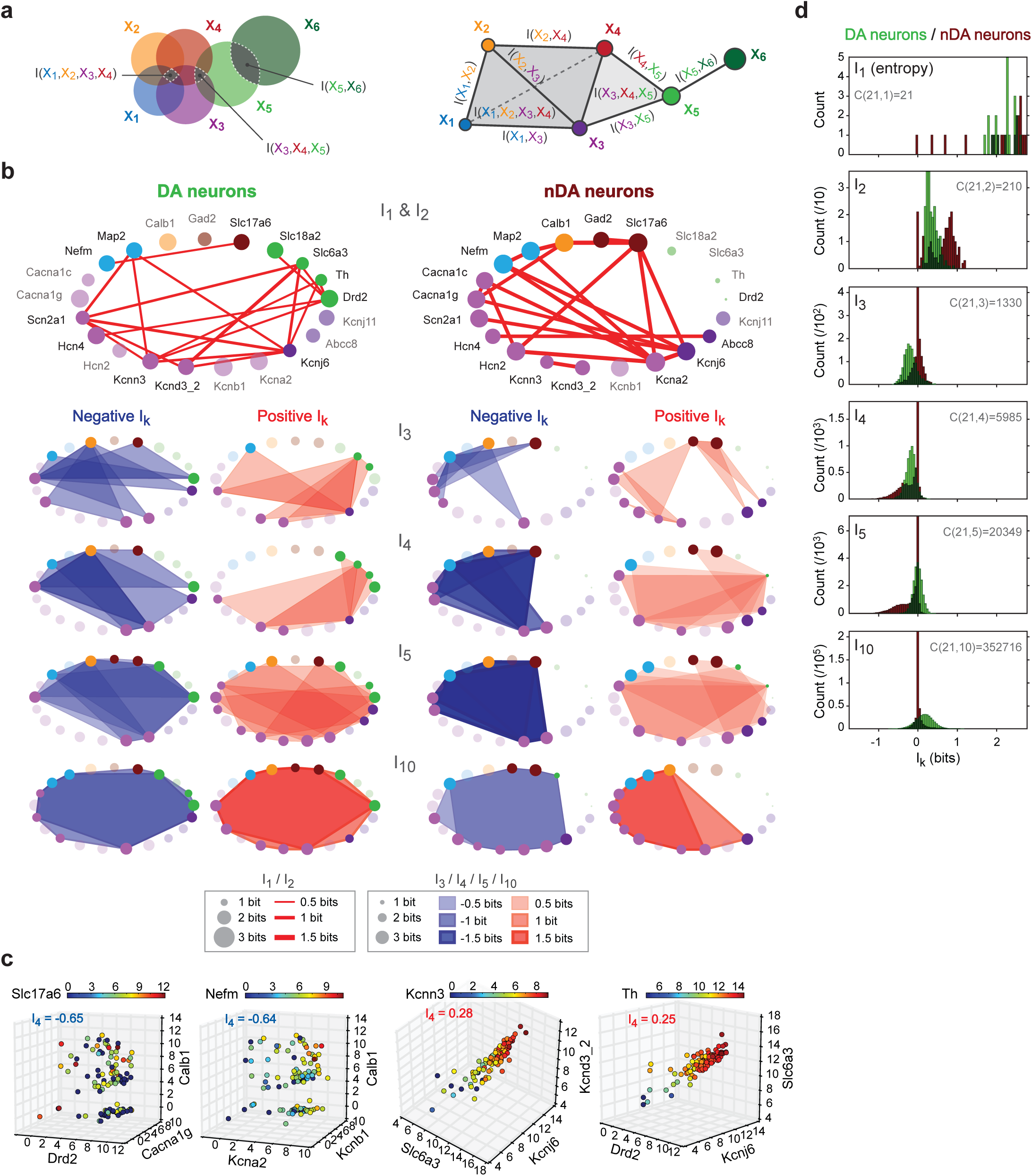
Mutual information analysis of gene expression levels reveals specific high-degree structures of transcriptomic profiles in midbrain DA and nDA neurons. **a**, left, Venn diagram illustrating a system of 6 random variables sharing mutual information at degree 2, 3 and 4. Right, same system represented on a simplicial complex. Each vertex represents a variable while the edges, faces and volumes represent joint-*n*-tuples of variables. **b**, scaffold representations of the most significant I_k_ values shared by pairs (I_2_, top row), triplets (I_3_, second row), quadruplets (I_4_, third row), quintuplets (I_5_, fourth row) and decuplets (I_10_, fifth row) of genes in DA (left column) and nDA neurons (right column). Circle diameters are scaled according to entropy value (I_1_). The red shapes (lines, triangles, quadrilaterals, etc) indicate positive I_k_ shared by genes while the blue shapes correspond to negative I_k_. Only the most significant values of I_k_ are displayed on each scaffold (20 for I_2_; 5 positive and 5 negative for I_3_, I_4_, I_5_; and 2 positive and 2 negative for I_10_). **c**, 4D-scatter plots representing the levels of expression of 2 quadruplets of genes sharing strong negative I_k_ (left-hand plots) and 2 quadruplets of genes sharing strong positive I_k_ (right-hand plots) in DA neurons. Please note that negative I_k_ is associated with a “clustering” or “heterogeneous” distribution of gene expression levels (left) while positive I_k_ is associated with a “co-varying” or “homogeneous” distribution of gene expression levels. **d**, histograms representing the distribution of I_k_ values for all the degrees presented in **b**. The total number of combinations C(*n*,*k*) for each degree (number of pairs for I_2_; number of triplets for I_3_, etc) is given in gray.

We applied I_k_ analysis to the gene expression levels measured in DA and nDA neurons for the 21 most relevant genes (**Figure 2**). The variability in expression of each gene X_i_ is quantified by the entropy H_1_(X_i_)=I_1_(X_i_) (**Supplementary methods**). Consistent with the expression profiles depicted in **Figure 1b**, the smallest and largest values of I_1_ were found for nDA neurons (**Figure 2b,d**). The genes sharing the strongest I_2_ values (**Figure 2b**) significantly overlapped with those sharing strong Pearson correlations (**Figure 1d**), in particular for DA neurons (*Th*/TH, *Slc6a3*/DAT, *Slc18a2*/VMAT2, *Drd2*/D2R, *Kcnj6*/GIRK2, *Kcnd3_2*/Kv4.3, *Kcnn3*/SK3, *Scn2a1*/Nav1.2). Nevertheless the precise patterns of I_2_-sharing genes were different, due to the fact that I_k_ also identifies non-linear dependences^**23**^. Interestingly, for *k* ≥ 3, the modules of genes sharing the strongest positive I_3_ and I_4_ displayed dense overlap with those sharing the strongest I_2_, while the groups of genes sharing the strongest negative I_3_ and I_4_ (*Cacna1g*/Cav3.1, *Calb1*/CB, *Drd2*/D2R, *Kcna2*/Kv1.2, *Kcnb1*/Kv2.1, *Kcnj11*/Kir6.2, *Nefm*/NEF3, *Slc17a6*/VGLUT2) had very little overlap with the strongly correlated (see **Figure 1d**) or strong I_2_-sharing genes (**Figure 2b**), especially for DA neurons. I_k_ was also calculated for superior degrees (5 to 21), and examples of the strongest positive and negative information modules are shown for I_5_ and I_10_ in **Figure 2b**. Consistent with the theoretical examples presented in **Supplementary Figure 6**, strong negative I_4_ was associated with clustering patterns of expression while strong positive I_4_ corresponded to co-varying patterns of expression (**Figure 2c**). In general, the distribution of I_k_ at each degree was found to be very different between DA and nDA neurons, with a predominance of independence (0 values) and strong negative values in nDA compared to DA neurons (**Figure 2d**).

In order to provide an exhaustive picture of the statistical dependences in both populations, we determined the information landscapes corresponding to the distribution of I_k_ values as a function of degree *k* (**Figure 3a**, **Supplementary Figure 7**). To help the reader understand this representation, two theoretical examples are given in **Supplementary Figure 7b**: for randomly equidistributed (independent) variables, I_1_ = log_2_(N), and I_2,…,n_ = 0; while for strictly redundant variables (*e.g.* correlation of 1), I_1,…,n_ = log_2_(N). The information landscapes of DA and nDA neurons were found to be very different from these two theoretical examples and from each other: in particular, the landscape of nDA neurons mainly comprised strong negative and 0 I_k_ values for k ≥ 3, suggesting that most *k*-tuples of genes are *k*-independent in these neurons. The prevalence of *k*-independence was found to be even stronger when the information landscape was computed for the 20 “less-relevant” genes in DA and nDA neurons (**Supplementary Figure 7c**). In contrast, the information landscape of DA neurons showed a predominance of negative I_k_ for *k* < 5 and predominance of positive I_k_ for *k* ≥ 5 (**Figure 3a**). Therefore, this analysis revealed a complex combinatorial structure of gene expression profiles in DA and nDA neurons, mixing independent, synergistic and redundant *k*-tuples of genes for *k* ≥ 3. In analogy with mean-field approximations, we also calculated the mean information for all degrees (**Figure 3a**, **Supplementary methods**). Due to the rather small number of cells analyzed and the inherent undersampling issue, the information landscapes computed here (especially the mean landscapes) should be intepreted with caution for *k* > 6 (DA) and *k* > 5 (nDA), even though maximal positive and negative I_k_ values are less sensitive to this limit (see **Supplementary methods**).

**Figure 3.**
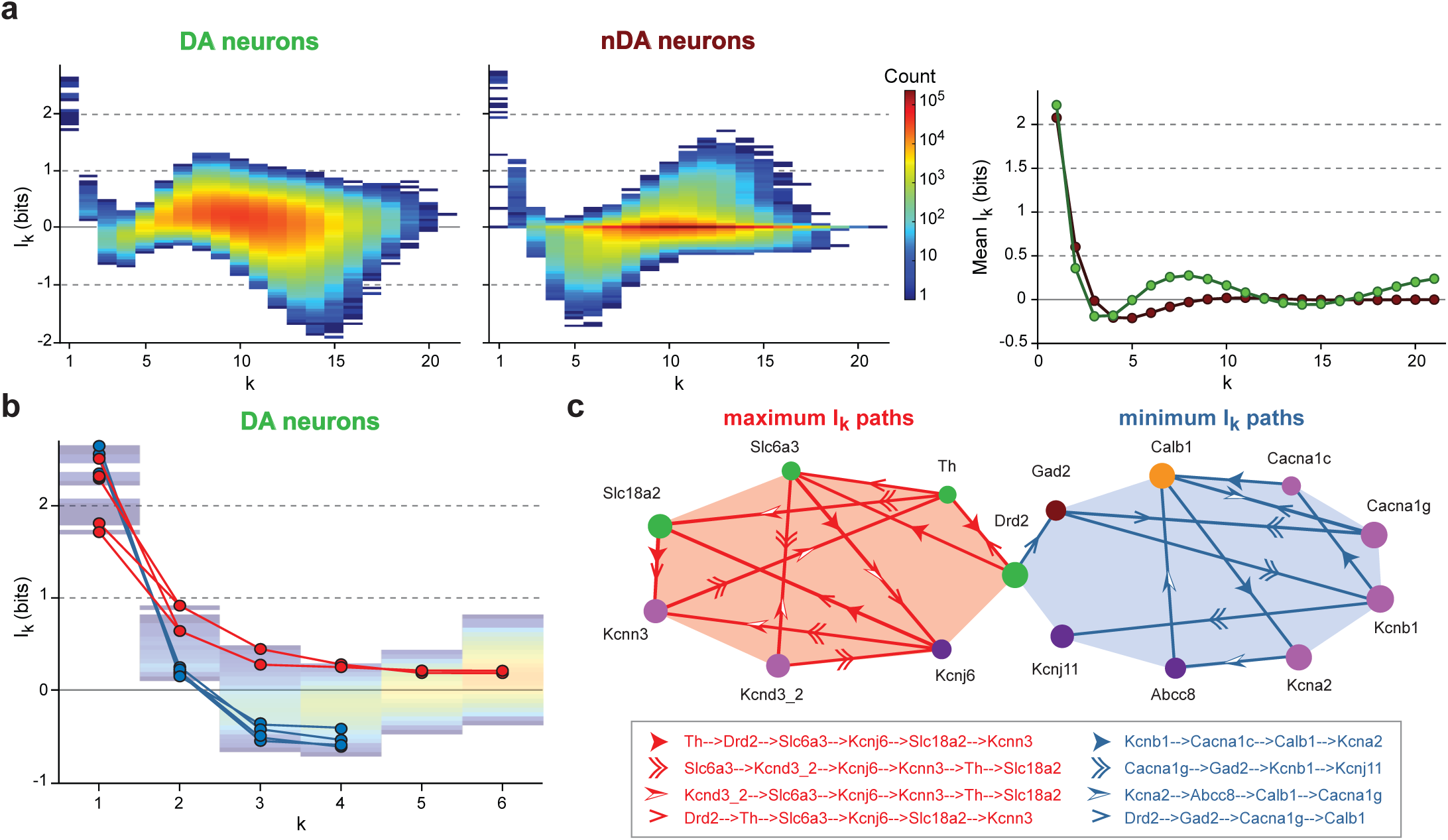
Conditional mutual information identifies stable modules of genes sharing strong statistical dependences. **a**, information landscapes representing the distribution of I_k_ values shared by the 21 genes presented in **Figure 2** as a function of degree *k* in DA (left) and nDA neurons (middle), and scatter plot representing mean I_k_ value (mean information landscape) as a function of degree for both populations (right). Color coding in the left and middle plots indicates the density of points for each I_k_ value. **b**, line and scatter plot illustrating the four maximum (red) and minimum (blue) information paths corresponding to stable information modules in DA neurons identified using conditional information computation. The paths have been superimposed to the total information landscape (transparent color coding) already shown in panel **a**. **c**, scaffold representation of the information modules corresponding to the maximum I_k_ paths (red) and minimum information paths delineated in panel **b**. Each path is identified with a specific arrowhead shape (see legend box).

The I_k_ analysis presented in **Figure 2** revealed that modules of strong positive or negative I_k_ could persist across degrees, but did not allow us to estimate the size of these gene modules. In order to quantify the stability of information modules and determine their size, we estimated the information flow over paths in the lattice of random variables in DA neurons (**Figure 3b-c, Supplementary Figure 8**). For a given information path, the first derivative with respect to the degree *k* is given by the conditional mutual information with a minus sign (**Supplementary methods**):

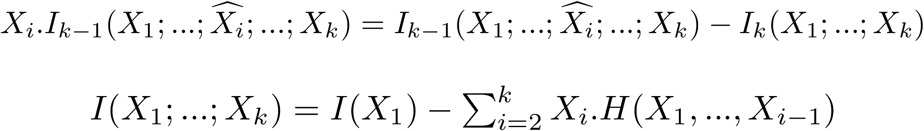

where ^ denotes the omission of X_i_ (the conditioning variable). X_i_.I_k-1_ stays positive (negative slope) if adding a variable X_i_ to the module increases the information while a negative X_i_.I_k-1_ (positive slope) indicates that adding a variable increases the uncertainty about the module. Therefore, reaching the first minima X_i_.I_k-1_ = 0 indicates that adding a variable stops being informationally relevant, and allows to define the degree for which information modules become unstable. In other words, the degree of the first minima gives a definitive assessment of the size of a gene module.

We characterized the paths that maximized mutual information (most informative modules) or that minimize mutual information (sequence of variables that segregate the most the whole set of variables), and that stay stable (**Supplementary methods**). **Figure 3b** presents the 4 longest paths of maximal and minimal information, which correspond to stable modules of degree 6 and 4, respectively. We then built the scaffold composed of the 4 maximal and minimal information paths (**Figure 3c**). All the genes involved in defining DA metabolism and signaling were found in the scaffold of maximal paths (*Th*/TH, *Slc6a3*/DAT, *Slc18a2*/VMAT2, *Drd2*/D2R), together with three ion channel genes (*Kcnj6*/GIRK2, *Kcnd3_2*/Kv4.3, *Kcnn3*/SK3), in keeping with the pairs, triplets and quadruplets of positive I_k_-sharing genes identified in **Figure 2b**. This finding brings new insights to our understanding of gene regulation in DA neurons. As shown in **Figure 2c**, the genes sharing strong positive I_k_ have co-varying profiles of expression, which is usually considered to indicate a co-regulation of expression^**4, 8**^. Therefore the positive information module determined using conditional mutual information (**Figure 3c**) should correspond to a group of genes co-targeted by the same regulatory factors. Several studies have demonstrated that the expression levels of *Th*/TH, *Slc6a3*/DAT, *Slc18a2*/VMAT2 and *Drd2*/D2R are indeed under the control of the same pair of transcription factors *Nurr1*/*Pitx3* ^**24**^ (**Supplementary Figure 9**). Our results are consistent with these observations, but moreover suggest that these four genes might be part of a larger transcriptional module (≥ 7 genes) that also includes genes defining the electrical phenotype of DA neurons (*Kcnj6*/GIRK2, *Kcnd3_2*/Kv4.3, *Kcnn3*/SK3). This also means that defining the neurotransmitter identity and the electrical phenotype of these neurons might be the product of a single transcriptional program, involving at least the *Nurr1* and *Pitx3* transcription factors (**Supplementary Figure 9**). Alternatively, this coupling between ion channel and DA metabolism genes might also reflect the documented activity-dependent regulation of DA-specific genes such as *Th*/TH, which has been shown to be sensitive to blockade of sodium (including Nav1.2) and potassium (including SK3) channel activity^**25**^.

On the other hand, the minimal information paths identified the 8 genes that best segregate midbrain DA neurons (**Figure 3c**), supporting the already documented diversity of this neuronal population^**12-14**^. The presence of *Abcc8*/SUR1*, Cacna1g*/Cav3.1, *Calb1*/CB*, Gad2*/GAD65*, Kcnj11*/Kir6.2 and *Drd2*/D2R is perfectly consistent with several studies linking the expression of these genes to specific subpopulations of SNc and VTA neurons ^**13, 26-29**^ (**Supplementary Figure 9**). Importantly, our analysis reveals that other genes, in particular the potassium channels *Kcna2*/Kv1.2 and *Kcnb1*/Kv2.1 might be used as markers of midbrain DA neuron subpopulations.

In summary, we showed that the topology of a high-dimensional dataset defined by the independence, and the simple (redundant) and complex (synergistic) statistical dependences at all degrees can be estimated using multivariate mutual information analysis (I_k_). Applied to sc-RTqPCR data, I_k_ analysis allowed us to simultaneously determine the size and identity of gene regulatory modules conserved across a cell population and the size and identity of gene modules underlying cell diversity (**Supplementary Figure 9**). Therefore, the specific complex combinatorial structure of genetic interactions (positive, negative, null) underlying the stability and diversity of a given cell type is described at once by the presented method. While applied here to transcriptomics data, I_k_ analysis could be applied to any type of high-dimensional data, within the limit of computational tractability.

## ACKNOWLEDGEMENTS

This work was funded by the French National Research Agency (ANR JCJC grant ROBUSTEX to J.M.G.; supporting S.T.), the European Research Council (ERC consolidator grant 616827 *CanaloHmics* to J.M.G.; supporting M.T.P., P.B. and M.L.), and the French Ministry of Research (doctoral fellowship to M.A.D.). We would like to thank Pr. E. Marder for helpful discussions on the manuscript.

## MATERIAL AND METHODS

### Acute midbrain slice preparation

Acute slices were prepared from P14–P23 TH-GFP mice (transgenic mice expressing GFP under the control of the tyrosine hydroxylase promoter) ^**30**^ of either sex. All experiments were performed according to the European and institutional guidelines for the care and use of laboratory animals (Council Directive 86/609/EEC and French National Research Council). Mice were anesthetized with isoflurane (Piramidal Healthcare Uk) and decapitated. The brain was immersed briefly in oxygenated ice-cold low calcium artificial cerebrospinal fluid (aCSF) containing the following (in mM): 125 NaCl, 25 NaHCO_3_, 2.5 KCl, 1.25 NaH_2_PO_4_, 0.5 CaCl_2_, 4 MgCl_2_, 25 glucose, pH 7.4, oxygenated with 95% O_2_ / 5% CO_2_ gas. The cortices were removed and then coronal midbrain slices (250 μm) were cut on a vibratome (Leica VT 1200S) in oxygenated ice-cold low calcium aCSF. Following 30–45 min incubation in 32°C oxygenated low calcium aCSF, the slices were incubated for at least 30 min in oxygenated aCSF (125 NaCl, 25 NaHCO_3_, 2.5 KCl, 1.25 NaH_2_PO_4_, 2 CaCl_2_, 2 MgCl_2_ and 25 glucose, pH 7.4, oxygenated with 95% O_2_ / 5% CO_2_ gas) at room temperature prior to electrophysiological recordings. Picrotoxin (100 μM, Sigma Aldrich, St. Louis, MO) and Kynurenate (2 mM, Sigma Aldrich) were bath-applied via continuous perfusion in aCSF to block inhibitory and excitatory synaptic activity, respectively.

### Cell dissociation and collection

Midbrain DA neurons were acutely dissociated following a modified version of the methods described in references ^**31**^ and ^**32**^. Regions containing the SNc, part of the VTA and SNr were excised from each coronal midbrain slice. The tissue was submitted to papain digestion (2.5 mg/ml and 5mM L-cysteine) for 15-20 min in oxygenated low calcium HEPES aCSF (containing 10 mM HEPES, pH adjusted to 7.4 with NaOH) at 35-37° C and subsequently rinsed in low-calcium HEPES aCSF supplemented with trypsin inhibitor and bovine serum albumin (1mg/ml). Single cells were isolated by gentle trituration with fire-polished Pasteur pipettes and plated on poly-L-Lysine-coated coverslips. Dissociated cells were maintained in culture in low calcium HEPES aCSF at 37° in 5% CO_2_ for at least 45 minutes. Coverslips were then placed in a cell chamber of a fluorescence microscope and continuously perfused with HEPES-aCSF. Cells were collected by aspiration into borosilicate glass pipettes mounted on a micromanipulator under visual control. Cell dissociation and collection were performed using RNA-protective technique and all solutions were prepared with RNase-free reagents when possible and filtered before use.

### Electrophysiology recordings, data acquisition and analysis

All recordings were performed as already described previously ^**33**^. Picrotoxin and Kynurenate were present for all recordings to prevent contamination of the intrinsic activity by spontaneous glutamatergic and GABAergic synaptic activity. Statistical analysis (performed according to data distribution) included: unpaired *t* test, Mann Whitney, paired *t* test with a p value <0.05 being considered statistically significant. Statistics were performed utilizing SigmaPlot 10.0 (Jandel Scientific, UK) and Prism 6 (GraphPad Software, Inc., La Jolla, CA).

### qPCR assays, specific retro-transcription and targeted amplification (RT-STA)

Pre-designed TaqMan assays (TaqMan^®^ Gene Expression Assays, Thermo Fisher Scientific) used in this study are listed in **Supplementary Table 1**. Assays were systematically selected to target the coding region and to cover all known splice variants. In the case of *Kcnd3* and *Kcnj6* genes, two different assays were used to detect all known splice variants. Excluding *Fos* (754 bp intron) and *Bdnf*, *Kcna2* and *Kcnj11* (both primers and probe within a single exon), assays spanning a large intron (>1000 bp) were chosen to avoid DNA amplification. *Gad1* primers and probe were designed according to Applied Biosystems criteria and MIQE recommendations ^**34**^. TaqMan^®^ assays were pooled (0.2x final concentration) and the preamplification step was validated using log serial dilutions of mouse brain total RNA (MBTR) ^**5, 6**^. The following thermal profile was applied: 50°C for 15 min, 95°C for 2 min and 22 cycles of amplification ^**35**^ (95°C for 15 s and 60°C for 4 min) following Fluidigm recommendations. For each assay, efficiency was estimated from the slope of the standard curve using the formula E= (10^(-1/slope)]^-1) x100. All assay efficiencies (89.4≤ E ≤100.4 %) are listed in **Supplementary Table 1.**

### Single-cell RTqPCR, data processing and analysis

Individual GFP and non-GFP neurons were harvested directly into 5 μl of 2x Reaction buffer (CellsDirect™ One-Step qRTPCR, Lifetech) and kept at −80°C until further processing. A reverse transcription followed by a specific targeted pre-amplification (RT-STA) was performed in the same tube (2.5 μl 0.2x assay pool; 0.5 μl SuperScript III) applying the same thermal profile described above. The pre-amplified products were treated with ExoSAPI (Affimetrix) and diluted 5-fold prior to analysis by qPCR using 96.96 Dynamic Arrays on a BioMark System (BioMark™ HD Fluidigm). Data were analyzed using Fluidigm Real-Time PCR Analysis software (Linear Baseline Correction Method and User detector Ct, Threshold Method). Two genes, *Kcnj6_c* and *Chat* were undetectable in all analyzed cells. Cells that had a Ct for *Hprt* above 21 were excluded from further analysis. After interplate calibration, all Ct values were converted into relative expression levels using the equation Log_2_Ex = Ct_LOD_ - Ct_(Assay)_ ^**36**^. LOD (limit of detection) was set to Ct=25 by calculating the theoretical Ct value for 1 single molecule in the Biomark system from two custom-designed oligonucleotides: *Slc17a6* and *Penk*. All data pre-processing was performed in Microsoft Excel (Microsoft, Redmond, USA). Heatmap and correlation maps (Pearson correlation coefficient values excluding zero values, p value <0.5, n >5) were generated in the R environment (R Core Team 2016) using gplots, heatmap3, Hmics and corrplot packages. Gene expression scatter plots and frequency distribution plots were created in SigmaPlot 10.0 (Jandel Scientific) and Prism 6 (GraphPad Software, Inc, La Jolla, CA). Figures were prepared using Adobe Illustrator CS6.

### Topological information data analysis

The present analysis is based on the information cohomology framework developed by Baudot and Bennequin^**3**^ and relies on theorems establishing uniquely the usual entropy (H_k_) and multivariate mutual information (I_k_) as the first class of cohomology and coboundaries respectively with finite (non-asymptotic) methods (see **Supplementary Methods** for more detail).

### Simplicial Information structures

The information functions are defined on the whole lattice of partitions of the probability simplex of atomic probabilities, providing the general random variable lattice of joint-variables. The application of this framework to data analysis is developed in the subcase of simplicial information homology, which consists in the exploration of the simplicial sublattice of “set of subsets” defined dually for joint and mutual (meet) monoid structures of random variables, and whose exploration follows binomial combinatorics with a complexity in *O*(2^n^). It allows an exhaustive estimation of the information structure, that is the joint-entropy H_k_ and the mutual information I_k_, on all degrees *k* and for every *k*-tuple of variables (gene expression levels), defined respectively by the following equations:

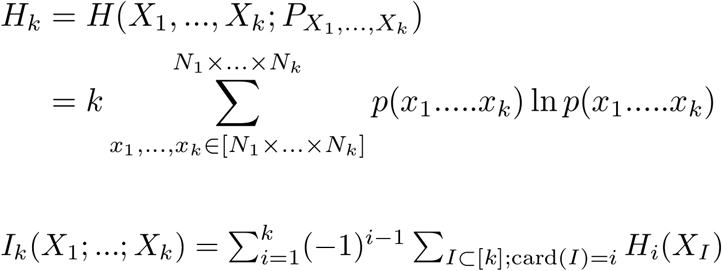

for a probability joint-distribution P_X1,…,Xk_ and joint-random variables (X_1_,…,X_k_) with alphabet [N_1_…N_k_] and k=-1/ln2, where *n* variables are mutually independent if and only if ∀ *k* ≤ *n*, I_k_=0. Due to the combinatorial complexity, in the current study H_k_ and I_k_ values were computed for *n*=21 (for *n*=21, the total number of information elements to estimate is 2 097 152).

The distributions of I_k_ and H_k_ for every degree *k* (corresponding to *k*-tuples of variables) were represented as I_k_ and H_k_ landscapes (**Supplementary Figure 7**). The landscapes are representations of the simplicial information structures where each element of the lattice is represented as a function of its corresponding value of entropy or mutual information, and quantify the variability-ramdomness and statistical dependencies at all degrees *k*, respectivey, from 1 to *n*. Mean landscapes were calculated by averaging I_k_ and H_k_ for each degree *k* over the number of *k*-tuples. The mean information landscape quantifies the average behavior of the whole structure. The mean information landscape (or path) is given by:

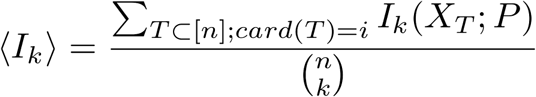

### Probability estimation

The probability estimation procedure is explained in **Supplementary Figure 5** for the simple case of two random variables (the expression levels of two genes). For each variable *X_j_*, we consider the space in the intervals [min *x_j_*, max *x_j_*] and divide it into *N_j_* boxes, *N* being the graining of the data. The empirical joint probability is estimated by box counting after a graining of the data space into *N_1_…N_k_* boxes (for *k*-tuple probability estimation). In the current study, a graining of *N_1_*=…=*N_k_*=8 was chosen as it provided a correct description of the distribution of the expressions levels (see **Supplementary Figure 8** for the influence of changing the graining on the identification of gene modules).

### Information paths

An information path IP_k_ or HP_k_ of degree *k* on I_k_ or a H_k_ landscape is defined as a sequence of elements of the lattice that begins at the leastest element of the lattice (the identity-constant “0”), travels along edges from element to element of increasing degree of the lattice and ends at the greatest element of the lattice of degree *k*. The first derivative of an IP_k_ path is minus the conditional mutual information. The (“non-Shannonian”) information inequalities ^**19**^, e.g. the negativity of conditional mutual information that quantifies the instability of the mutual information along the path, are then equivalent to the existence of local minima on such paths (see **Supplementary methods**). The critical dimension of an IP_k_ path is the degree of its first minima. A positive information path is an information path from 0 to a given I_k_ corresponding to a given *k*-tuple of variables such that I_k_<I_k-1_<…<I_1_. We call the marginal component of a path I_1_ a self-information energy and the interacting compoenents functions I_k_, *k* > 1, a free information energy. A maximal positive information path is a positive information path of maximal length: it ends at a minima of the free information energy function. In the current study, the length of maximal positive information paths was considered to indicate the size of a stable information module. The set of all these paths defines uniquely the minimum free information complex (see **Supplementary methods**). In simple terms, this complex is the homological formulation of the minimum energy principle with potentially many local and degenerate minima. The set of all paths of degree *k* is in one-to-one correspondence with the symmetric group Sk and hence untractable computationally (complexity in *O*(*k*!)). In order to bypass this issue, we used a fast local algorithm that selects at each element of degree *k* of an IP path the positive information path with maximal or minimal I_k+1_ value or stops whenever X_k_.I_k+1_ ≤ 0 and rank those paths by their length.

### Robustness of the method

To estimate the degree after which the sample size *m* becomes limiting and biases our estimations, the undersampling regime was quantified by the degree *k*_u_ beyond which a significant proportion (10%) of the H_k_ values get close to log_2_(*m*). Using these criteria, with log_2_(111)=6.79 and log_2_(37)=5.21, the *k*_u_ obtained for DA neurons was 6 and 5 for nDA neurons, and I_k_ and H_k_ values beyond these degrees should be interpreted with caution (**Supplementary methods**). It must be noted however that this limit is calculated on the average H_k_, whose value is mainly determined by non-relevant independent *k*-tuples. The biologically relevant statistical dependences correspond to extrema in the raw landscape (minimal H_k_ and maximal or minimal I_k_) and therefore are less affected by this sampling problem. In order to evaluate the robustness of our results to sample size (*m*) and graining value (*N*), we calculated the maximal positive paths obtained for DA neurons for smaller samples (*m* = 28, 56, 84, taken fully arbitrarily among 111) and smaller (*N*=4, 6) or larger (*N*=10, 12) graining values (**Supplementary Figure 8**). The information paths of maximal length were found to be relatively robust to variations in *N* and *m*, even though, as expected, m=28 yielded significantly different paths. For most *N* and *m* combinations, the main genes identified in **Figure 3c** were also present in the maximal information paths, including in particular the DA metabolism/signaling genes and the two ion channel genes *Kcnd3*/Kv4.3 and *Kcnn3*/SK3. Concerning the statistical significance of the results, I_2_ functions are Kullback-Leibler divergences ^**37**^ and estimate the divergence from 2-independence. Their generalization to arbitrary degree *k* (I_k_) can be interpreted as a statistical significance of a test, here against the null hypothesis of *k*-independence I_k_=0. Our analysis, based on the ranking of the I_k_ for every *k*, considered only the 5 maximal (positive) and 5 minimal (negative) values of I_k_, which are the 5 most significantly dependent positive and negative I_k_-sharing k-tuples (for *k* > 2).

### Computation and algorithm

The Information Topology open source program, written in Python, is available on Github depository. It allows to compute the information landscapes, paths, and minimum free energy complex, which encode and represent directly all the usual equalities, inequalities, and functions of information theory (as justified at length in **Supplementary methods**), and all the structures of the statistical dependences within a given set of empirical measures (up to the approximations, computational tractability and finite size biases, see previous sections). It can be run on a regular personal computer up to *k* = *n* = 21 random-variables in reasonable time (3 hours), and provide new tools for pattern detection, dimensionality reduction, ranking and clustering based on a unified homological and informational theory.

